# Orthogonal microfluidic approaches reveal force-enhanced migration of bacterial populations on surfaces

**DOI:** 10.64898/2026.09.10.750177

**Authors:** Piyush Sharma, Michael J. Marcheschi, Alexander M. Shuppara, Joseph E. Sanfilippo

## Abstract

Host-generated flow is expected to oppose bacterial migration by sweeping cells downstream, yet some bacteria can migrate upstream against the direction of flow. However, it is unclear how the magnitude of shear force influences upstream bacterial migration. Here, we use microfluidics to examine upstream migration of the human pathogen *Pseudomonas aeruginosa* under host-relevant shear forces (0.1–10 pN). By independently varying flow rate and solution viscosity, we discover that increasing shear force counterintuitively enhances population-level upstream migration. Single-cell tracking reveals that force-enhanced migration is driven by an increase in twitching motility speed. Using a microfluidic-based trigonometry approach, we link the increase in speed to the cell-surface angle. At low shear forces, type IV pilus retraction generates a torque that tilts cells toward a vertical orientation, geometrically constraining forward movement and decreasing twitching speed. In contrast, higher shear forces push cells toward a horizontal orientation, increasing twitching speed. Collectively, our results reveal how shear force can enhance bacterial motility and upstream migration, providing a framework to understand how host shear forces may promote the spread of bacterial infections.

## Introduction

For several centuries, scientists have observed bacterial motility in laboratory conditions (1, 2). However, bacteria in nature must contend with mechanical forces (3–5) that are often absent in simplified experiments. Fluid flow is ubiquitous in natural and host environments, but it is technically challenging to create experimental systems where flow is precisely controlled. To overcome this challenge, collaborative efforts between microbiologists and engineers have used microfluidic-based approaches (5–7). By simultaneously applying flow and observing bacterial cells, it has become clear that flow has a major impact on bacterial behavior (5). For example, flow suppresses quorum sensing (8, 9), enhances bacterial growth under nutrient limitation (10, 11), potentiates oxidative stress (12–14), and improves antibiotic effectiveness (15, 16). As incorporating flow into experiments has led to unexpected and exciting results, there is a clear need to use microfluidic approaches to study bacteria in flowing environments.

Flow exerts a shear force on surface-attached cells. In a microfluidic device, cells experience a shear force that is dependent on flow rate and solution viscosity (17–19). To test if flow-sensitive behaviors are triggered by shear force, researchers should perform experiments where flow rate is held constant while solution viscosity is altered. This strategy has helped define several key flow-sensitive bacterial processes. For example, changing viscosity revealed that the *Escherichia coli* adhesin FimH has catch bond properties (20), and *Pseudomonas aeruginosa* surface residence time is enhanced by shear force (18). Alternatively, changing viscosity revealed that flow-sensitive gene expression in *P. aeruginosa* is force-independent (17). By combining the power of microfluidics with viscosity experiments, there is a great opportunity to provide mechanistic understanding of flow-sensitive bacterial responses.

Many bacterial species migrate on surfaces via twitching motility (21–26). Twitching is carried out by extension and retraction of bacterial appendages known as type IV pili (22, 27). Type IV pili are comprised of repeating PilA monomers that typically extend out 0.5–2 µm from the cell pole (18, 24, 28) and can adhere to surfaces (29). Pilus retraction is rapid (∼0.5 µm/s) (27, 28), is powered by ATP (30), and generates approximately 10–100 pN of force (30–34). For surface-attached cells, successive extension, adhesion, and retraction events can pull cells forward (27). Mutant cells lacking functional type IV pili display reduced virulence (35, 36), suggesting that twitching plays an important role during infection. Microfluidic studies of twitching have revealed that flow can orient cells to twitch upstream (26, 37). As bacterial species from different ecological niches exhibit flow-induced upstream twitching (38, 39), the ability to twitch into the direction of flow appears to be an important bacterial adaptation.

However, the relationship between shear force and upstream twitching remains largely unexplored. Here, we use orthogonal microfluidic approaches to explore how shear force impacts upstream twitching of *P. aeruginosa*. During upstream twitching in flow, type IV pili pull cells forward and tilt cells off the surface. By independently modulating flow rate and solution viscosity, we demonstrate that higher shear forces counteract pilus-mediated cell tilting and push cells toward the surface. Surprisingly, we discover that higher shear forces lead to faster twitching, which contributes to better upstream migration of the population. Our counterintuitive discovery, that shear force enhances upstream migration, highlights the power of studying bacteria with microfluidics and has the potential to change how we think about bacterial pathogens in flowing host environments.

## Results

To investigate how shear flow impacts surface migration of bacterial populations, we imaged *P. aeruginosa* PA14 cells in microfluidic devices. For our first experiment, we built a microfluidic device with a branched design (Figure S1), which allowed us to create a reservoir of cells at the end of a cell-free channel (Figure 1A). We used a syringe pump to precisely flow media from left to right. As *P. aeruginosa* cells can use type IV pili to twitch upstream (37, 38), we hypothesized that cells would move from right to left and migrate up the empty channel.

**Figure 1:**
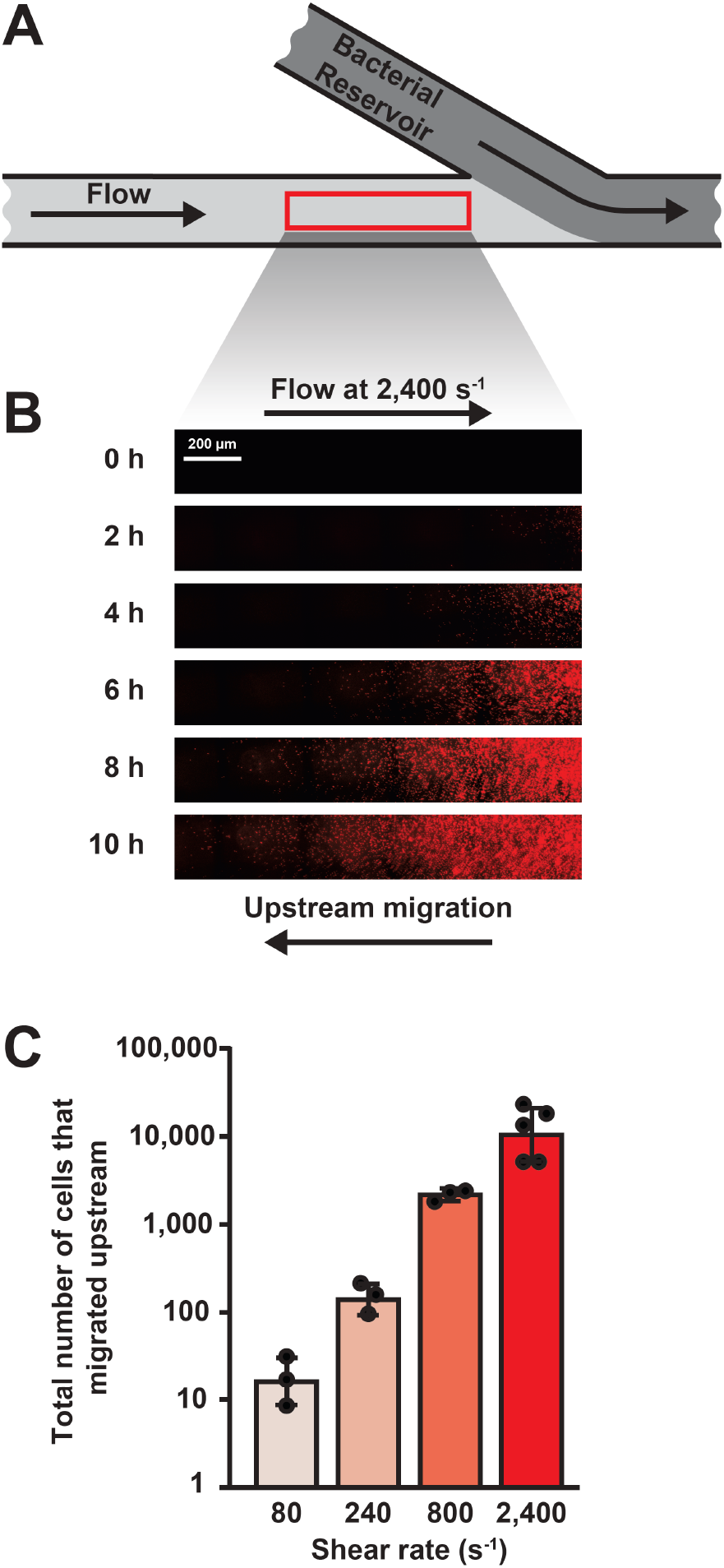
Shear flow enhances upstream migration of *P. aeruginosa* populations. **(A)** Branched channel microfluidic setup used to observe upstream surface migration. Cells were loaded through a side branch creating a bacterial reservoir (dark gray). Cells migrated upstream into the direction of flow into the cell-free main channel (light gray). **(B)** *P. aeruginosa* cells migrating upstream into constant flow with a shear rate of 2,400 s^-1^. Flow was directed from left to right, and cells migrated upstream from right to left. Lookup tables were independently optimized for each panel to improve bacterial population visualization. **(C)** Quantification of upstream surface migration at varied shear rates. Error bars represent SD of at least three biological replicates. Statistical significance was determined using Welch’s one-way ANOVA followed by Dunnett’s T3 test on log_10_ transformed data. All four experimental conditions were statistically different from one another (*p* < 0.05).

Shear rate describes the flow intensity and is calculated using flow rate and channel dimensions. Consistent with our hypothesis, cells exposed to a shear rate of 2,400 s^-1^ successfully migrated into the upstream portion of the channel over a 10-hour period (Figure 1B). To test if type IV pili were required, we repeated this experiment with a Δ*pilA* mutant that does not make pili. Δ*pilA* cells were unable to migrate into the upstream region (Figure S2), confirming that type IV pili are required for upstream migration. Thus, our custom-fabricated microfluidic device provided a useful system to study pilus-mediated upstream migration.

How does flow intensity impact upstream migration? We hypothesized that increasing flow would inhibit upstream migration. To test our hypothesis, we repeated our upstream migration experiment at a range of shear rates. We chose shear rates of 80, 240, 800, and 2,400 s^-1^ as they are commonly found in parts of the human body, such as the bloodstream (40, 41), urinary tract (42, 43), and lungs (44). When we exposed cells to a shear rate of 80 s^-1^, we observed very little upstream migration (Figure 1C). Surprisingly, cells exposed to higher shear rates exhibited a stepwise increase in migration (Figure 1C). In fact, cells exposed to a shear rate of 2,400 s^-1^ were the most successful (Figure 1C), steadily moving upstream throughout the 10-hour experiment (Figure 1B). Together, these experiments refuted our original hypothesis, established that increasing flow enhances upstream migration, and led us to search for the mechanism that underlies this counterintuitive behavior.

As upstream migration requires type IV pili, we reasoned that individual *P. aeruginosa* cells were twitching upstream in our devices. Using single-cell tracking in a straight microfluidic channel (Figure S1), we observed that ∼87% of cells twitch on surfaces when exposed to a shear rate of 800 s^-1^ (Figure S3). Of the cells that twitch at 800 s^-1^, we observed that ∼86% move in the upstream direction (Figure 2A, 2B). To more precisely quantify twitching motility, we tracked individual cells over a 30-minute period. We noticed that cells moved mostly straight in one direction with turns along the way (Figure 2A). Based on this observation, we quantified the displacement of cells over a 30-minute period (Figure 2C). Our tracking revealed that cells exposed to a shear rate of 800 s^-1^ had an average displacement of ∼24 µm (Figure 2C). These results establish how individual cells twitch in flow and allow us to investigate how flow intensity modulates twitching motility.

**Figure 2:**
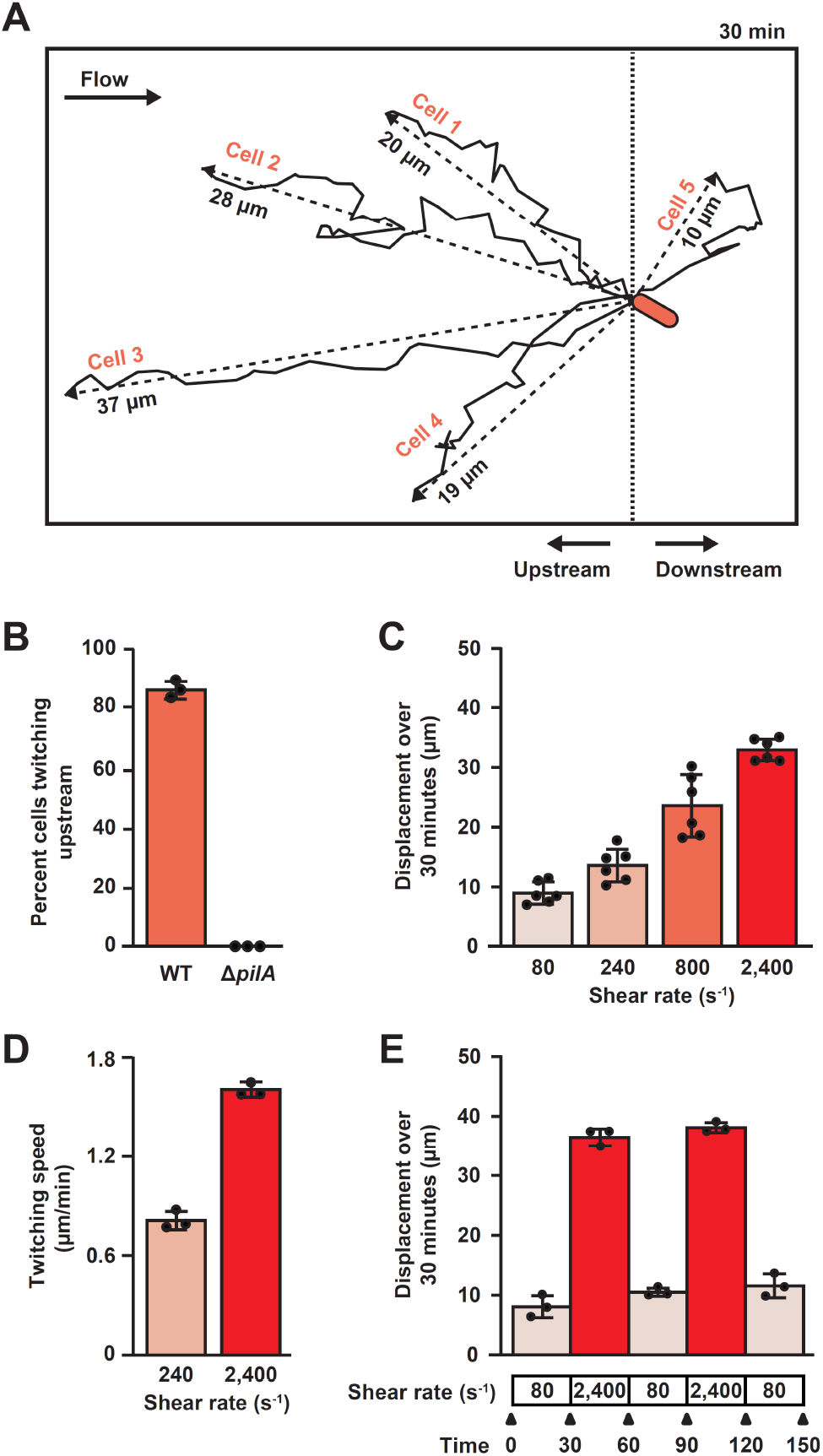
Shear flow enhances *P. aeruginosa* upstream twitching speed. **(A)** Five representative cell tracks on the surface over 30 minutes at a shear rate of 800 s^-1^. Displacement of each cell is shown via dashed lines, and distance moved by each cell is shown via solid lines. **(B)** Percentage of twitching WT and Δ*pilA* mutant cells that moved upstream at a shear rate of 800 s^-1^. Upstream was defined relative to the starting position of each cell. **(C)** Twitching displacement of WT cells at varied shear rates over 30 minutes. Displacement was defined as the distance between the start and end point for each cell. Statistical significance was determined using Welch’s one-way ANOVA followed by Dunnett’s T3 test. All four experimental conditions were statistically different from one another (*p* < 0.05). **(D)** Twitching speed of WT cells at varied shear rates. Statistical significance was determined using Student’s t-test (*p* < 0.01). **(E)** Twitching displacement of WT cells that were exposed to changing shear rates over time. Displacement repeatedly increased at high shear rate and repeatedly decreased at low shear rate. Error bars represent SD of at least three biological replicates. Each biological replicate represents an average of at least 30 cells.

How does flow intensity impact twitching motility? Based on our upstream migration results, we hypothesized that increasing flow would increase twitching. To test our hypothesis, we subjected *P. aeruginosa* cells to a range of shear rates and quantified twitching. We observed that ∼73% of cells exposed to a shear rate of 80 s^-1^ twitched and ∼93% of cells exposed to a shear rate of 2,400 s^-1^ twitched (Figure S3). Also, we observed that cells exposed to higher flow were slightly more biased toward moving upstream (Figure S4). Strikingly, when we quantified how far cells twitched in 30 minutes, we learned that cells exposed to higher shear rates exhibited a stepwise increase in twitching displacement (Figures 2C). In fact, cells exposed to a shear rate of 2,400 s^-1^ twitched 3.5 times farther than cells exposed to a shear rate of 80 s^-1^ (Figure 2C). Similarly, cells exposed to a shear rate of 2,400 s^-1^ twitched twice as fast as cells exposed to a shear rate of 240 s^-1^ (Figure 2D). Thus, flow can enhance twitching speed of *P. aeruginosa*.

We wondered if the ability of flow to enhance twitching was reversible. To test for reversibility, we tracked twitching of *P. aeruginosa* cells that were exposed to alternating 30-minute treatments of low flow (80 s^-1^) and high flow (2,400 s^-1^). The experiment started at a shear rate of 80 s^-1^, and cells had a low twitching displacement (Figure 2E). After we increased flow to 2,400 s^-1^, twitching displacement was approximately 4 times higher (Figure 2E). After we decreased flow to 80 s^-1^, twitching displacement returned to lower levels (Figure 2E). Twitching displacement increased and decreased again after additional alternating treatments, leading us to conclude that the ability of flow to modulate twitching is rapid and reversible. Collectively, our microfluidic experiments establish that flow enhances surface migration of *P. aeruginosa* populations by increasing twitching motility of individual cells.

While quantifying twitching, we noticed that wildtype *P. aeruginosa* cells were typically tilted off the surface at an angle (Figure 3A). We hypothesized that the cell-surface angle is controlled by type IV pili. In support of our hypothesis, we observed that Δ*pilA* mutant cells typically lay flat (Figure 3A). To calculate the cell-surface angle for individual cells, we used a trigonometry approach that considered the observed cell length (when a cell is tilted) and the full cell length (when a cell is pushed down by flow) (Figure S6). For example, we calculated that a cell with an observed length of 1.3 µm and a full length of 3.0 µm had a cell-surface angle of 64° (Figure S6). Using this approach, we determined that wildtype cells exposed to a shear rate of 800 s^-1^ had an average cell-surface angle of ∼40° and Δ*pilA* cells had an average cell-surface angle of ∼3° (Figure 3B). Based on these results, we conclude that *P. aeruginosa* cells tilt themselves off the surface in a pilus-dependent manner.

**Figure 3:**
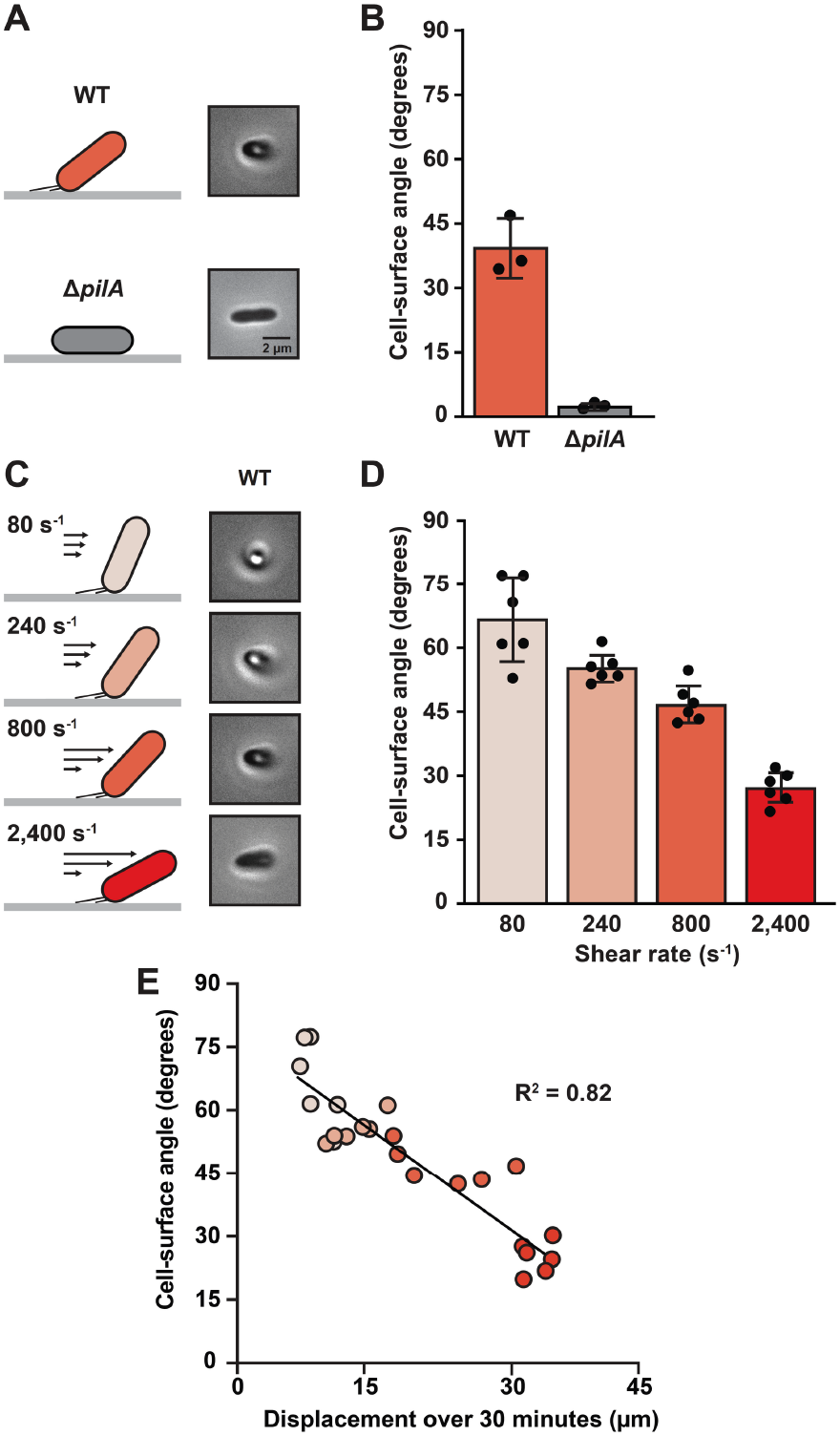
Type IV pili and shear flow set the *P. aeruginosa* cell-surface angle. **(A)** Cell schematics alongside phase-contrast images of WT and Δ*pilA* mutant cells exposed to a shear rate of 800 s^-1^. WT cells tilt off the surface in a pilus-dependent manner. **(B)** Quantification of the cell-surface angle for WT and Δ*pilA* mutant cells at a shear rate of 800 s^-1^. Cell-surface angle was calculated with a trigonometry approach explained in the methods. Statistical significance was determined using Student’s t-test (*p* < 0.05) **(C)** Cell schematics alongside phase-contrast images of WT cells exposed to varied shear rates. Cells are pushed closer to the surface at higher shear flow. **(D)** Quantification of the cell-surface angle of WT cells exposed to varied shear rates. Error bars represent SD of at least three biological replicates. Each biological replicate represents an average of at least 30 cells. Statistical significance was determined using Welch’s one-way ANOVA followed by Dunnett’s T3 test. All conditions except 80 s^-1^ versus 240 s^-1^ were statistically different from one another (*p* < 0.05). **(E)** Cell-surface angle correlates with twitching displacement. Individual data points represent average cell-surface angle (shown in 3D) and average twitching displacement (shown in 2C). The solid line indicates best-fit linear regression (*R^2^* = 0.82).

How does flow increase twitching speed? We propose a model where flow decreases the cell-surface angle, which then leads to an increase in twitching speed (Figure 2D). To examine this relationship, we used our trigonometry approach to calculate the cell-surface angle of wildtype cells at a range of shear rates. Consistent with our model, cells exposed to increasing flow had a stepwise reduction in their cell-surface angle (Figures 3C, 3D, S7).

Furthermore, we observed a strong negative correlation between cell-surface angle and twitching (Figure 3E). As cells with lower angles twitched farther than cells with higher angles (Figure 3E), we conclude that flow increases twitching by decreasing the cell-surface angle.

How does flow decrease the cell-surface angle? As flow imparts a shear force on surface-attached cells, we hypothesized that shear force decreases the cell-surface angle by pushing cells toward the surface. In our microfluidic channels, the shear force that cells experience is proportional to shear rate and solution viscosity (Figure 4A). Thus, if flow is pushing cells toward the surface, the cell-surface angle should decrease at both a higher shear rate and a higher solution viscosity. As cell-surface angle is impacted by shear rate (Figure 3D), we explicitly tested the impact of shear force by using the viscous agent Ficoll. When shear rate was held constant and viscosity was increased 10-fold (via addition of 15% Ficoll), cells had an average cell-surface angle of ∼31° (Figure 4B). Consistent with our hypothesis, when cells were exposed to flow with a shear rate of 240 s^-1^ without any change in solution viscosity, the average cell-surface angle was ∼57° (Figure 4B). Based on these results, we conclude that shear force decreases the cell-surface angle by pushing cells toward the surface.

**Figure 4:**
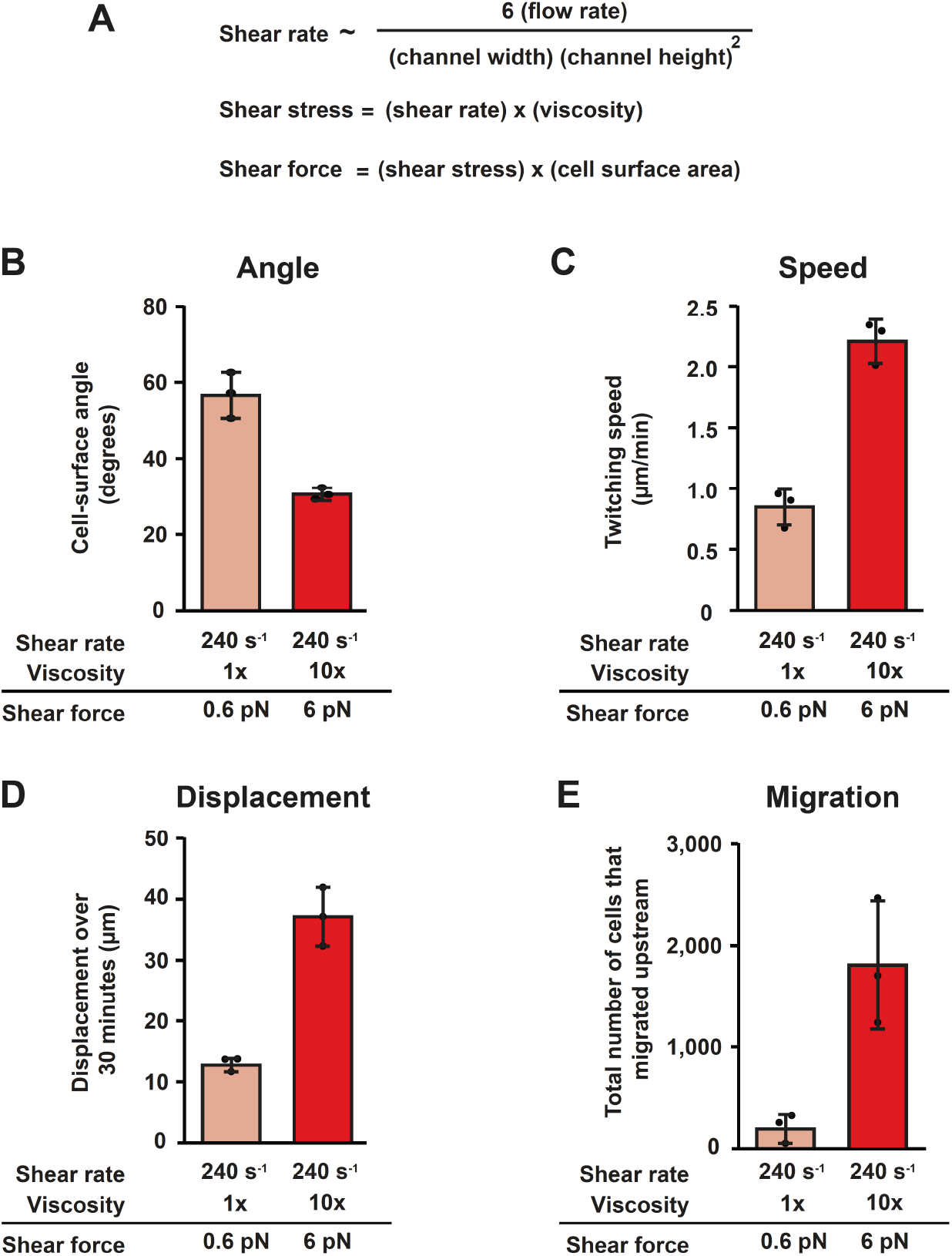
Increasing viscosity reveals that shear force mediates *P. aeruginosa* surface behaviors in flow. **(A)** Shear rate is dependent on flow rate, channel width, and channel height. Shear stress is proportional to shear rate and solution viscosity. Shear force is equal to shear stress times cell surface area. **(B)** Quantification of cell-surface angle at a shear rate of 240 s^-1^ with either no Ficoll (1x viscosity) or 15% Ficoll (10x viscosity). Each biological replicate represents an average of at least 30 cells. **(C)** Quantification of twitching speed at a shear rate of 240 s^-1^ with either no Ficoll or 15% Ficoll. Each biological replicate represents an average of at least 30 cells. **(D)** Quantification of twitching displacement at a shear rate of 240 s^-1^ with either no Ficoll or 15% Ficoll. Each biological replicate represents an average of at least 30 cells. **(E)** Quantification of population-level upstream surface migration at a shear rate of 240 s^-1^ with either no Ficoll or 15% Ficoll. Error bars represent SD of three biological replicates. All comparisons between 1x and 10x viscosity conditions were determined to be statistically significant using Student’s t-test (*p* < 0.05).

As shear force pushes cells toward the surface, we hypothesized that shear force increases twitching. To test our hypothesis, we altered solution viscosity (while holding shear rate constant) and measured twitching of individual cells. In support of our hypothesis, we found that increasing solution viscosity increased twitching speed (Figure 4C) and twitching displacement (Figure 4D). As shear force increases twitching, we predicted that shear force also enhances upstream surface migration. As predicted, we found that populations exposed to flow with increased solution viscosity were more successful at migrating upstream (Figure 4E). Thus, our results establish that shear force pushes cells toward the surface, leading to an increase in twitching speed, which ultimately enhances upstream migration of *P. aeruginosa* populations.

Collectively, our orthogonal microfluidic approaches reveal the counterintuitive relationship between shear force and bacterial migration, suggesting that bacterial pathogens can co-opt host mechanical forces to their benefit.

## Discussion

How does shear force impact surface migration of bacterial populations? Intuitively, shear force could inhibit upstream migration by sweeping bacterial cells downstream, with higher shear leading to even greater inhibition. Here, we provide evidence challenging this intuition. Using a combination of microfluidic approaches, we demonstrate that higher shear forces can instead promote the upstream migration of the human pathogen *P. aeruginosa*. Additionally, we characterize the biophysical mechanism underlying shear-enhanced upstream migration. In low shear regimes, pilus retraction generates a torque on cells, tilting them into a more vertical orientation (Figure 5A). In contrast, higher shear counteracts pilus retraction, pushing cells into a more horizontal orientation (Figure 5A). Cells in a horizontal orientation move faster, as they travel farther with each pilus retraction event (Figure 5A, 5B). Over time, shear-enhanced twitching promotes population-level upstream migration (Figure 5C). Thus, rather than simply opposing surface migration, higher shear forces can change cell-surface geometry in a manner that drives bacterial populations farther upstream.

**Figure 5:**
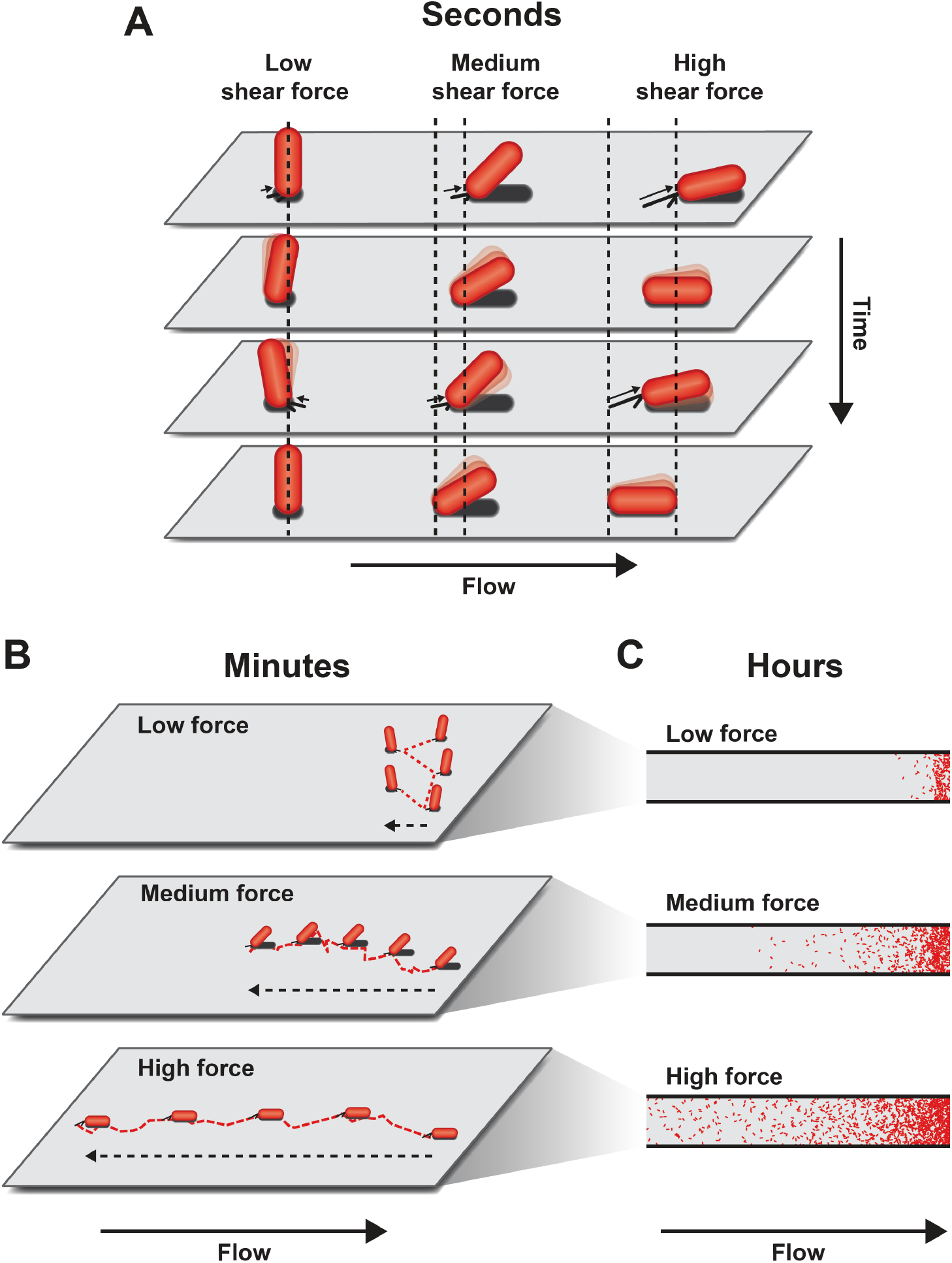
Shear force tips cells over, increases twitching motility, and enhances population-level migration. **(A)** Cells exposed to lower shear forces are more vertical and have less upstream movement during twitching. Cells exposed to higher shear forces are more horizontal and have more upstream movement during twitching. **(B)** Over minutes, cells exposed to higher shear forces get farther upstream than cells exposed to lower shear forces. **(C)** Over hours, bacterial populations exposed to higher shear forces migrate farther than populations exposed to lower shear forces.

What advantage do bacteria gain by twitching on surfaces? At first glance, twitching motility appears slow compared to faster modes of motility, such as flagellar-based swimming (2). However, many distantly related species twitch on surfaces (21, 24, 25, 45), suggesting that twitching provides an advantage in surface-attached contexts. While swimming allows bacteria to relocate over long distances in liquid, twitching allows bacteria to reposition themselves locally after landing on a surface. Although motility and adhesion are often framed as competing processes, twitching allows cells to be motile without leaving the surface. This combination of motility and adhesion is a particularly advantageous strategy in flowing environments, where swimming cells risk being swept downstream. We therefore postulate that twitching provides bacteria with a distinct advantage in flow, allowing cells to reposition themselves locally while remaining attached to the surface.

Upstream twitching has been observed in bacterial species from diverse ecological niches, including the human pathogen *P. aeruginosa* (24), the plant pathogen *Xylella fastidiosa* (26), and the thermophilic bacterium *Thermus thermophilus* (23). Although some reports have classified upstream twitching as rheotaxis, we have chosen not to use that term because there is no direct evidence that active sensing is involved. In the reported cases, upstream twitching is carried out by type IV pili that are polarly located on rod-shaped cells (23, 26, 28). The most parsimonious explanation is that pili serve as surface anchor points, flow physically reorients cells downstream of pili, and repeated pilus extension and retraction events pull cells upstream. Similarly, we propose that force-enhanced upstream twitching can be explained without invoking active sensing. Our results indicate that higher shear forces push cells toward a more horizontal cell-surface geometry, allowing cells to travel farther on the surface (Figure 3) (46). Together, our results lead us to propose that the interplay between external shear force, pilus retraction, and cell-surface geometry underlies the mechanism by which increasing shear enhances upstream twitching.

How do host mechanical forces impact surface migration? Physically confining cells, increasing flow, or increasing viscosity might be expected to restrict surface migration. However, as these mechanical changes all promote a more horizontal cell-surface geometry, our results indicate that they can instead enhance surface migration. Physical confinement between an agar pad and a solid surface facilitates twitching (47), likely because mechanical force promotes a more horizontal cell orientation. Flow intensities found in the bloodstream (40, 41) and urinary tract (42, 43) promote a more horizontal cell orientation (Figure 3) and enhance twitching (Figure 2), indicating that flow-enhanced twitching may occur during infection. Similarly, increasing solution viscosity across a physiologically relevant range (48, 49) promotes a more horizontal cell orientation (Figure 4B) and increases twitching (Figure 4C, 4D), suggesting that twitching may be more effective in viscous fluids. Collectively, our microfluidic experiments reveal that host-relevant mechanical forces can reorient cells with respect to the surface and counterintuitively enhance surface migration of bacterial populations.

## Acknowledgements

We thank Iota Chen, Anuradha Sharma, Evan Johnson, Will Wightkin, Matthias Koch, Josh Shaevitz, and Katja Taute for helpful discussions and comments on the manuscript.

## Contributions

P.S., M.J.M, A.M.S., and J.E.S. designed research. P.S. and M.J.M. performed research. P.S., M.J.M., and J.E.S. analyzed data. P.S., A.M.S., and J.E.S. designed the figures. P.S. and J.E.S. wrote the paper.

## Supplementary Information for

### Materials and Methods

#### Strains and growth conditions

The bacterial strains used in this paper are described in Table S1. *P. aeruginosa* PA14 cultures were grown in liquid LB in a roller drum at 37°C and on LB plates (1.5% Difco Agar). LB medium was prepared using premade Miller LB Broth (BD Scientific) using standard LB preparation protocols. Bacterial cultures were grown to mid-log with an approximate OD of 0.5.

#### Fabrication of microfluidic devices

Microfluidic devices were made as previously described (12). Briefly, microfluidic devices were created using soft lithography techniques. Photomasks were designed using Adobe Illustrator and printed through Artnet Pro Inc. Photomask patterns were embedded onto 100 mm silicon wafers (University Wafer) that were then spin-coated with SU-8 3050 photoresist (KAYAKLI Advanced Materials). Microfluidic chips were made with polydimethylsiloxane (PDMS) and plasma treated to bond to 60 mm x 36 mm x 0.16 mm glass coverslips (Ted Pella Inc.).

Migration experiments used branched channels with a branch channel dimension of 500 µm wide x 50 µm tall x 0.25 cm long and main channel dimension of 500 µm wide x 50 µm tall x 4.25 cm long (Figure S1). Twitching and cell-surface angle experiments were performed in straight microfluidic channels with channel dimensions of 500 µm wide x 50 µm tall x 2.0 cm long (Figure S1).

#### *P. aeruginosa* in microfluidic devices

For twitching and cell-surface angle experiments, microfluidic devices were loaded with bacteria as previously described (12). Briefly, bacterial cultures were loaded into the channels with a pipette. Cells were allowed to settle for ∼10 minutes before flow was turned on. All experiments were carried out at approximately 24°C with mid-log cultures of bacterial cells. Device inlets were attached to a length of polyethylene tubing, ID 0.38 mm x OD 1.09 mm (BD Intramedic Polyethylene) that was sheathed over a 26-gauge x 1/2” hypodermic needle (Air-tite Products) that was affixed on a syringe (BD). Device outlets were connected to another length of polyethylene tubing that dispensed the flowing media into a waste container. The inlet syringes were filled with LB and were mounted on a syringe pump (KD Scientific Legato 210) that was set to flow rates between 1–30 µL/min (corresponding to shear forces between 0.2–6 pN).

For migration experiments, a syringe filled with LB was fastened to a hypodermic needle. Polyethylene tubing was fastened to the end of the needle and tubing was inserted to the inlet of the main channel. A second length of polyethylene tubing was connected to the outlet and dispensed the flowing media into a waste container. The inlet syringe was mounted on a syringe pump, and flow rate was set to 3 µL/min. Media filled the main channel and branched channel. A sealed length of polyethylene tubing was inserted into the inlet of the branch channel to prevent media from flowing out.

After the sealed polyethylene tubing was inserted into the branch inlet, a 4 mL mid-log culture of WT *P. aeruginosa* cells constitutively expressing mCherry was concentrated to 1 mL using a centrifuge (Eppendorf Centrifuge 5420) set to 4,000 rpm for 4 minutes. Cells were drawn into a syringe via a single-use needle (Air Tite Disposable Needle, 16-gauge x 3”). The single-use needle was discarded, and the syringe was connected to a 26-gauge x 1/2” hypodermic needle. Polyethylene tubing was attached to the needle. The tubing was inserted into the branch inlet while flow continued in the main channel. The filled syringe was mounted on a second syringe pump. The pump was turned on for 5 minutes at a flow rate of 1.5 µL/min. After 5 minutes, the pump was turned off, and the syringe remained attached to the inlet via polyethylene tubing. A reservoir of *P. aeruginosa* cells was created at the rightmost end of the microfluidic device (Figure 1A).

#### Phase-contrast and fluorescence microscopy

Images were collected on a Nikon Eclipse Ti-2 microscope with the NIS Elements interface. Images were taken with a Nikon 40x Plan Apo Ph2 0.95 NA objective, a Nikon 100x Plan Apo Oil Ph3 1.45 NA objective, a Hamamatsu Orca-Flash 4.0 LT camera, and a Lumencor Sola Light Engine LED light source. Twitching and cell-surface angle experiment images were collected using the Nikon 100x Plan Apo Oil Ph3 1.45 NA objective with 1.5x zoom. Upstream migration experiment images were collected using the Nikon 40x Plan Apo Ph2 0.95 NA objective.

#### Shear rate and shear force calculations

Shear rate and shear force were calculated as previously described (18). Briefly, the wall shear rate experienced in a microfluidic device was calculated using the following equation:

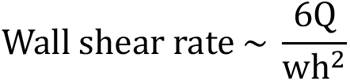

where Q is the flow rate, w is the channel width, and h is the channel height. As shown in Figure 4A, shear stress was calculated to be the product of shear rate and solution viscosity. Shear force was calculated to be the product of shear stress and the surface area of a cell, which was estimated to be 2.5 μm^2^ (17), as shown in Figure 4A.

#### Increasing solution viscosity

Solution viscosity was increased 10-fold by adding 15% Ficoll (MP Biomedicals) to LB medium. 15% Ficoll has been previously shown to increase solution viscosity 10-fold (17).

#### Quantification of migration in branched channel microfluidic

Phase-contrast and fluorescent images were captured in 2-hour intervals over 10 hours to quantify upstream surface migration. To quantify upstream surface migration, images were stitched together to create a large image with dimensions of 7 mm x 1 mm. Quantification of total number of cells migrated upstream was carried out at the 10-hour interval using ImageJ. The main channel was divided into twenty 100 µm x 500 µm segments. At shear rates of 80 s^-1^, 240 s^-1^, and 800 s^-1^, cells were manually counted in each segment and total number of cells was quantified by adding up each segment.

At the shear rate of 2,400 s^-1^, cell density was very high. As a result, manually counting the total number of cells was technically challenging. To overcome this challenge, the main channel was divided into twenty 100 µm x 500 µm segments. The number of cells in each segment was estimated by dividing each 100 µm x 500 µm segment into 100 µm x 25 µm boxes from top of the channel to the bottom. Total cells were manually quantified in the topmost 100 µm x 25 µm box. Subsequent boxes in a single 100 µm x 500 µm segment that had similar cell density were counted, and total number of cells was calculated for each box. The rest of the cells in the 100 µm x 500 µm segment were manually quantified. This process was repeated for each 100 µm x 500 µm segment and total number of cells was calculated.

#### Quantification of twitching in straight channel microfluidic devices

For our experiments, we defined twitching as total cell movement of at least 3 µm, which is approximately one cell body length. To quantify twitching displacement and twitching speed, phase-contrast images were captured every 30 seconds for 30 minutes. Captured videos were analyzed in ImageJ. To measure displacement, 30 cells that stayed on the surface for the entire 30 minutes were randomly selected. Cell displacement was quantified by measuring the distance between the cell’s starting point and ending point.

To measure twitching speed, 30 cells that stayed on the surface for the entire 30 minutes were randomly selected. For each cell, the twitching speed was quantified by measuring the distance moved by the cell between each frame (30-second intervals). We chose 30-second intervals because it is visually challenging to differentiate twitching and Brownian motion when using shorter intervals.

#### Quantification of cell-surface angle

To quantify cell-surface angle, phase-contrast images were captured every 30 seconds over 30 minutes. At the end of 30 minutes, flow was increased to a shear rate of 8,000 s^-1^ and images were captured for another 2 minutes. Using ImageJ, 30 random cells were chosen. Cell length was measured twice for the same cell: once at the experimental flow condition and once at 100 µL/min (which is a shear rate of 8,000 s^-1^).

To calculate the cell-surface angle, we used the following trigonometry equation:

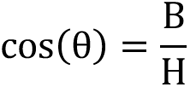

where *θ* = cell-surface angle, B = cell length under the experimental flow condition and H = cell length at 8,000 s^-1^.

To account for measurement error when cells are completely vertical, the following correction was employed:

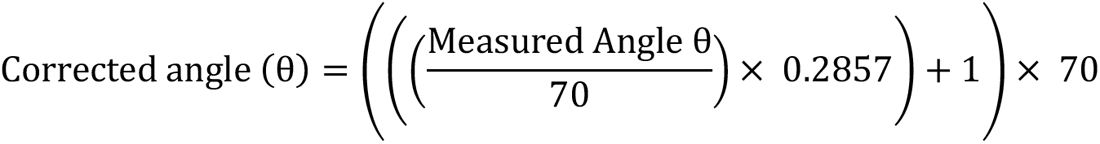

where 70 degrees is the highest measured angle recorded using the trigonometry equation.

---

**Table S1:** Strains used in this study.

| *P. aeruginosa* strain | Description | Source |
| --- | --- | --- |
| PA14 | wildtype, clinical isolate from burn wound | (50) |
| JS12 | attB::[plac-mCherry FRT] | (17) |
| JS176 | $\Delta pilA::FRT$ | (18) |

**Figure S1:**
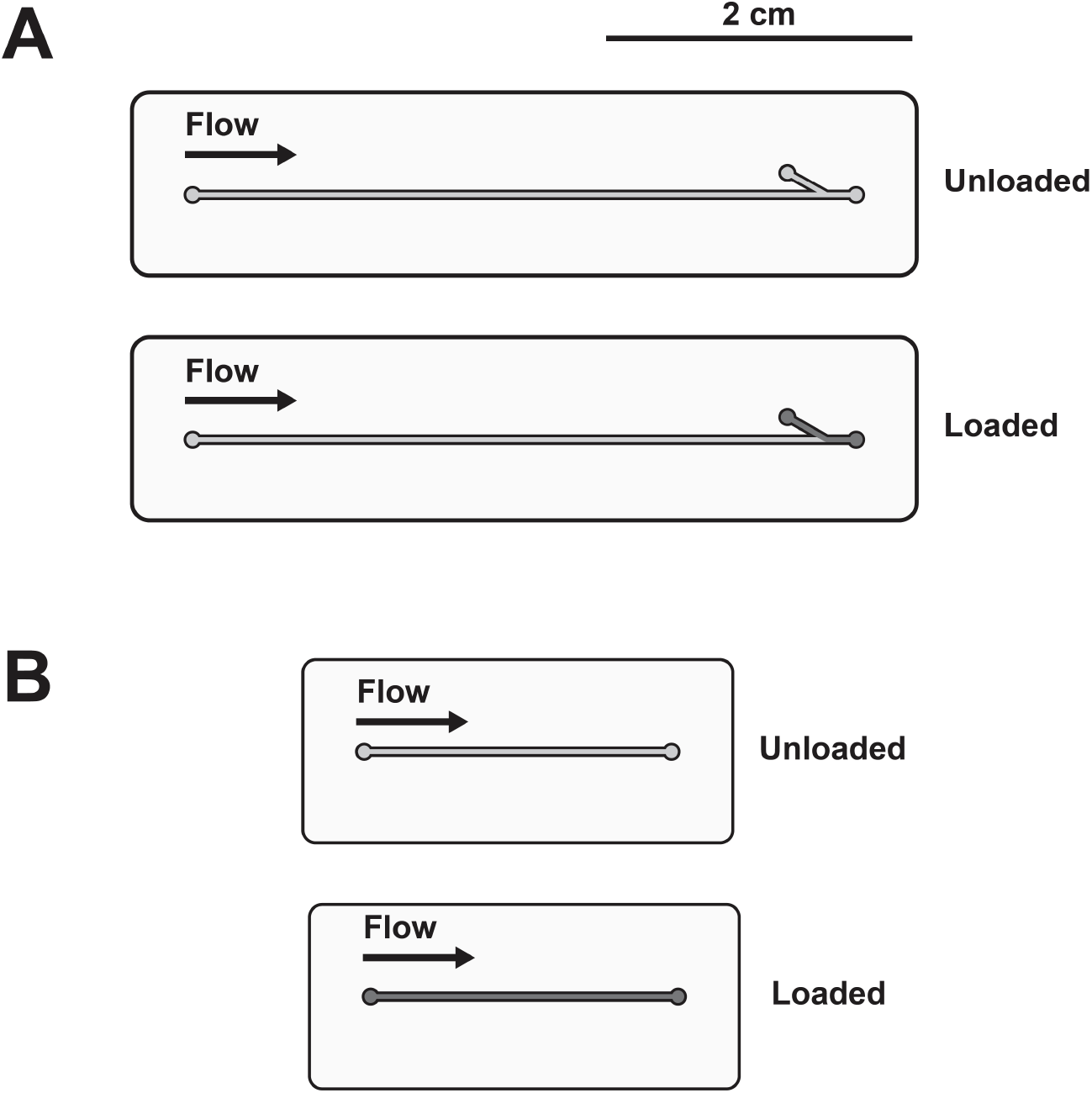
Schematics of microfluidic setups used throughout this study **(A)** Branched channel microfluidic used for upstream surface migration experiments. The microfluidic was comprised of a cell-free main branch (500 µm wide x 50 µm tall x 4.25 cm long) and a side branch (500 µm wide x 50 µm tall x 0.25 cm long). *P. aeruginosa* cells were loaded through the side branch creating a bacterial reservoir on the right-hand side (dark grey). Cells migrated upstream into the cell-free main channel (light grey). **(B)** 2-cm straight channel used for twitching and cell-surface angle experiments. *P. aeruginosa* cells were loaded into the straight channel (dark grey). Each microfluidic schematic is drawn to scale.

**Figure S2:**
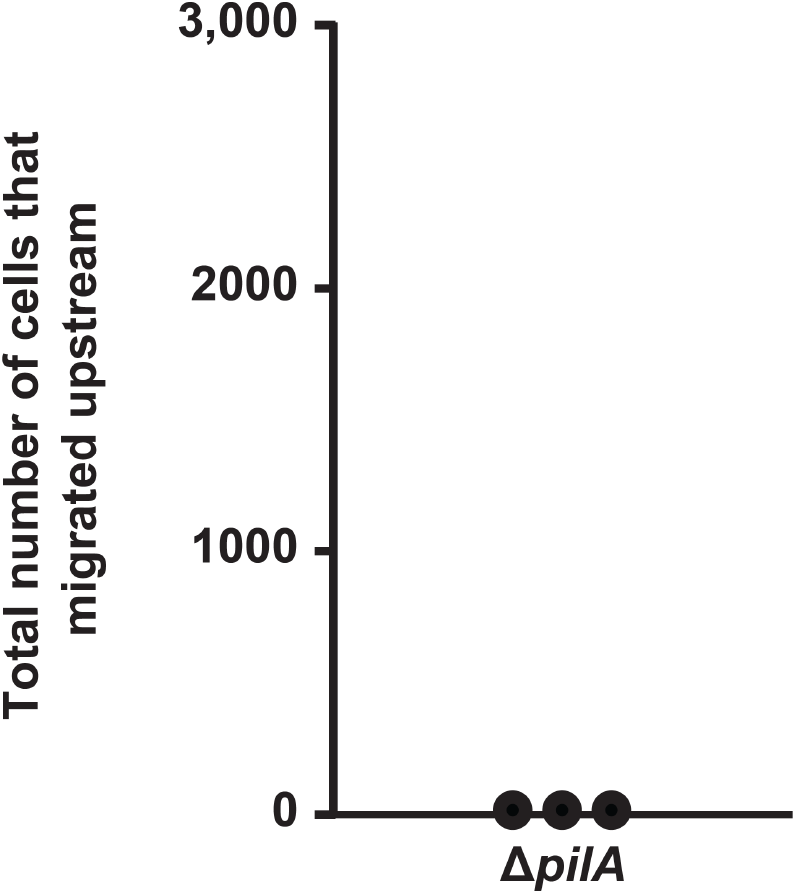
Type IV pili are required for upstream migration Δ*pilA* mutant cells do not migrate upstream. Cells were observed for 10 hours and provided with constant flow at a shear rate of 800 s^-1^. Each biological replicate represents an average of 30 cells.

**Figure S3:**
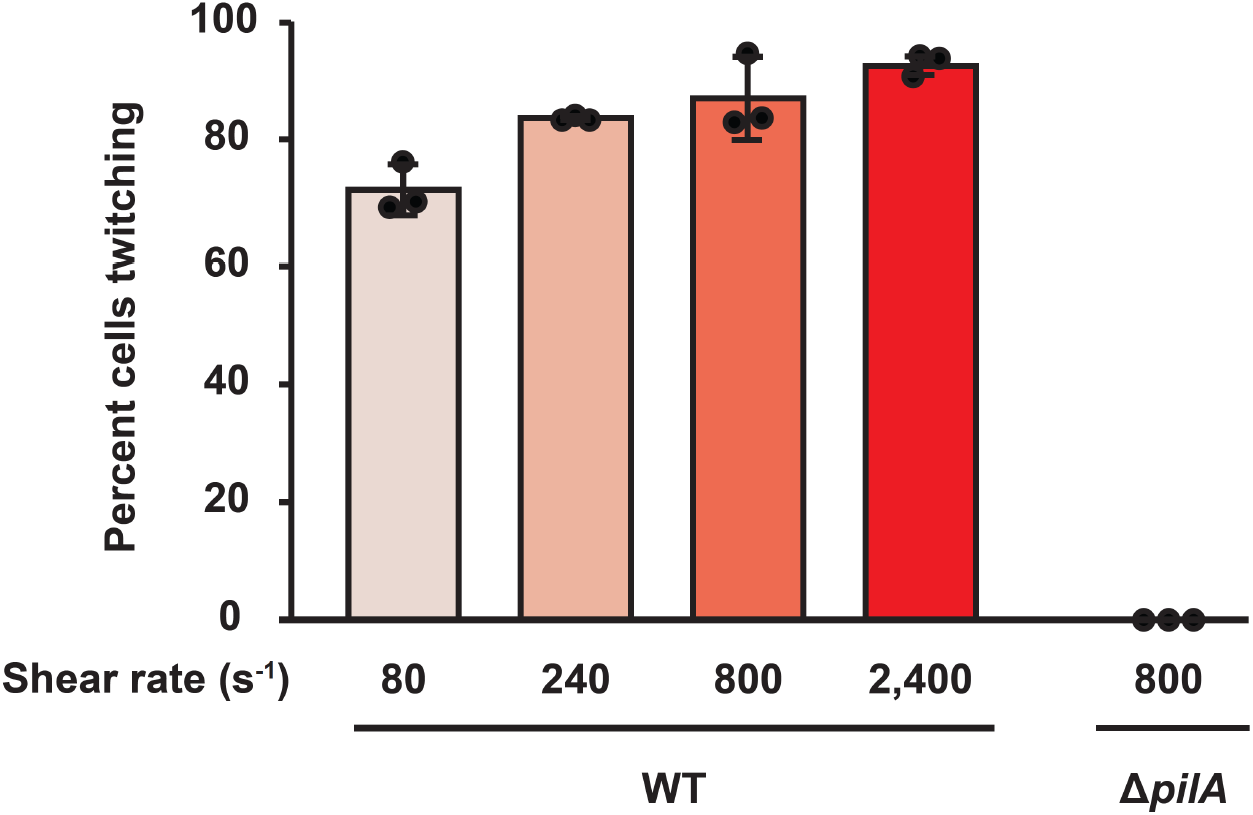
Increasing flow results in a modest increase in the percentage of cells undergoing type IV pili-dependent twitching Percentage of WT and Δ*pilA* mutant cells that twitched at varied shear rates over 30 minutes. Twitching was defined as a cell moving greater than one body length over 30 minutes. Error bars represent SD of three biological replicates. Each biological replicate represents an average of at least 30 cells. Statistical significance between WT conditions was determined using Welch’s one-way ANOVA followed by Dunnett’s T3 test. 80 s^-1^ versus 2,400 s^-1^ and 240 s^-1^ versus 2,400 s^-1^ were statistically different from one another (*p* < 0.05). All other WT conditions were not statistically different.

**Figure S4:**
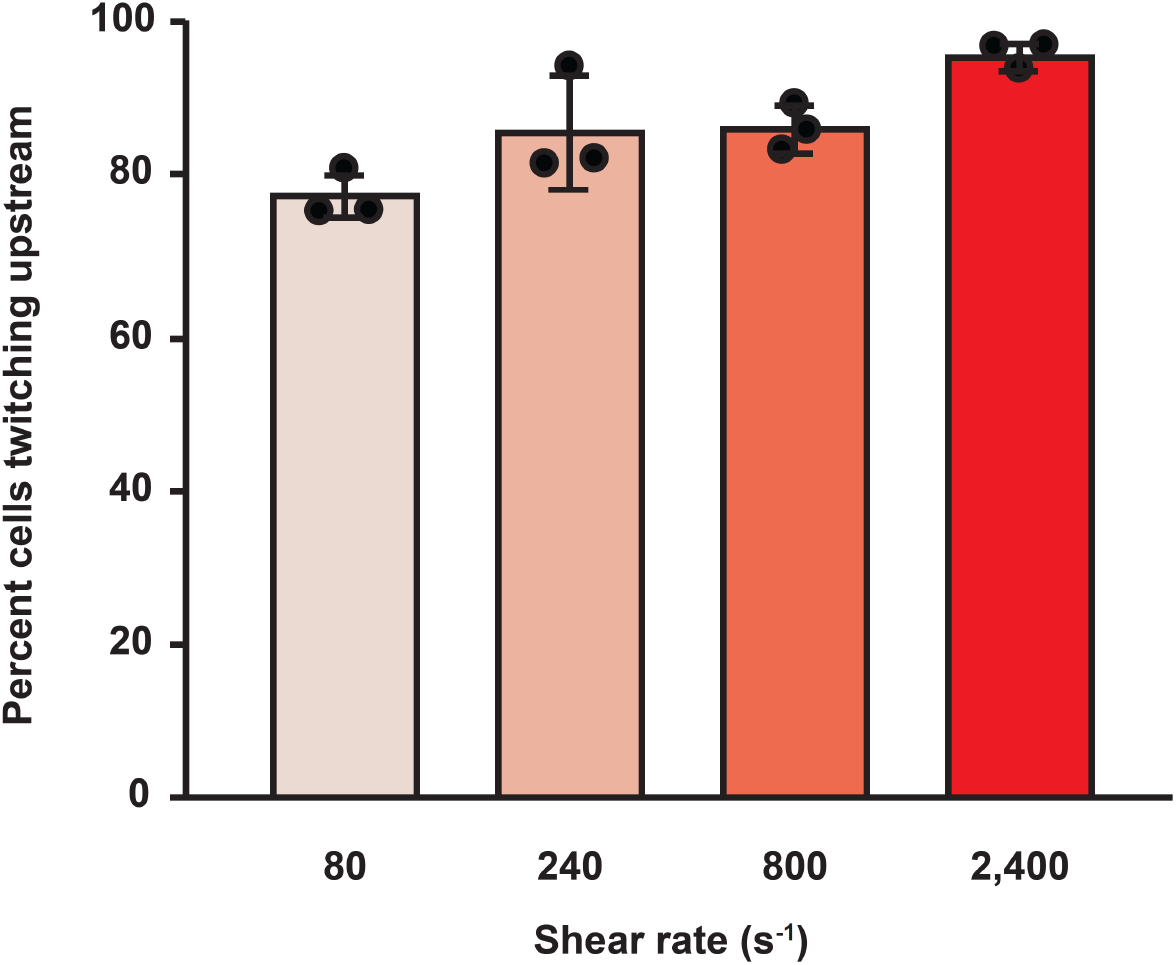
Most *P. aeruginosa* PA14 cells twitch upstream at shear rates between 80 s^-1^ and 2,400 s^-1^ Percentage of twitching WT cells that moved upstream at varied shear rates over 30 minutes. Upstream was defined relative to the starting position of each cell. Error bars represent SD of three biological replicates. Each biological replicate represents an average of at least 30 cells. Statistical significance was determined using Welch’s one-way ANOVA followed by Dunnett’s T3 test. 80 s^-1^ versus 2,400 s^-1^ were statistically different from one another (*p* < 0.01). All other conditions were not statistically different.

**Figure S5:**
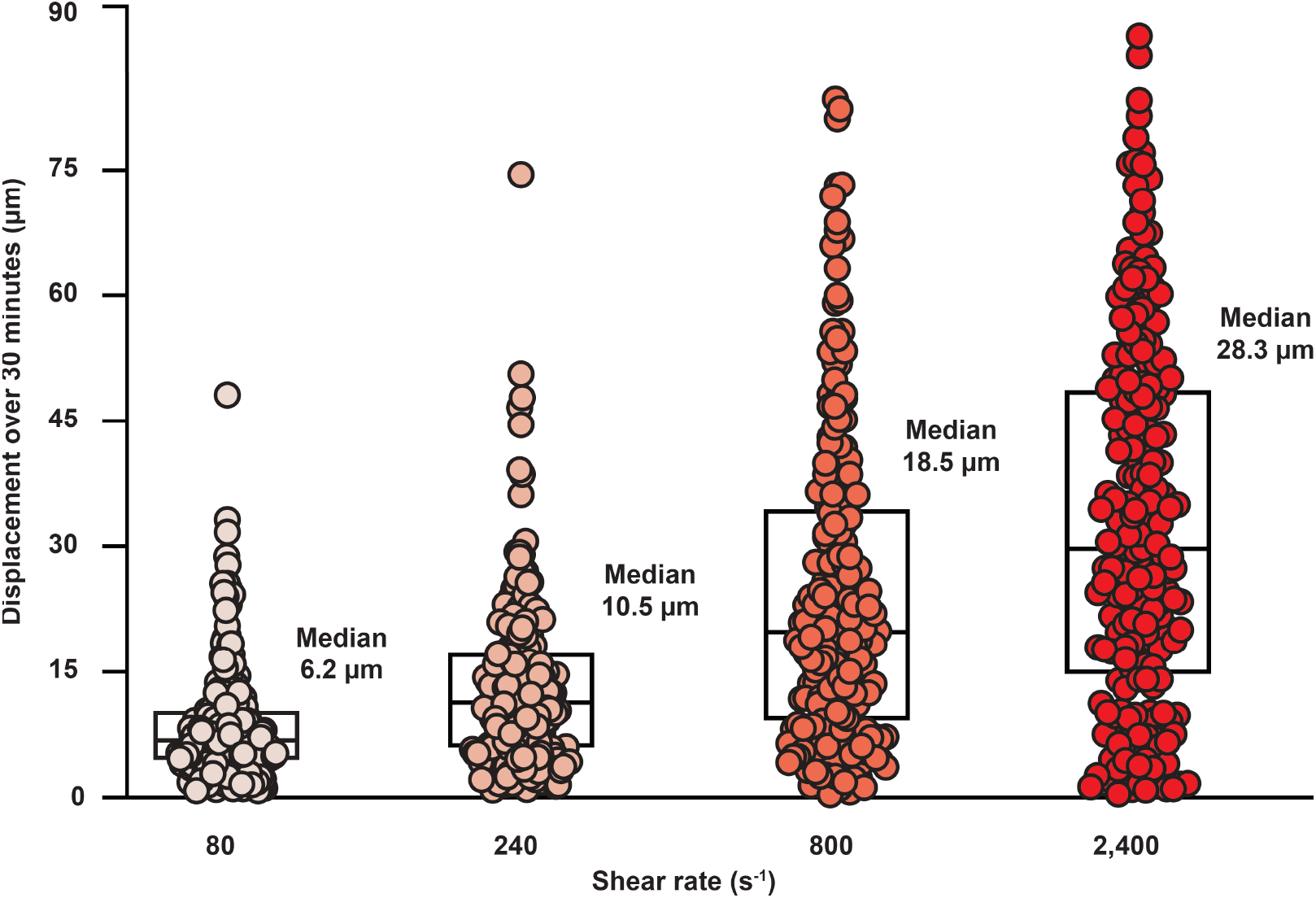
Increasing flow results in increased twitching displacement of individual cells Twitching displacement of individual WT cells at varied shear rates over 30 minutes. Displacement was defined as the distance between the start and end point for each cell. N = 180 cells for each condition. Statistical significance was determined using Welch’s one-way ANOVA followed by Dunnett’s T3 test. All four experimental conditions were statistically different from one another (*p* < 0.001). Boxplot represents the 25th percentile, median, and 75th percentile for each condition.

**Figure S6:**
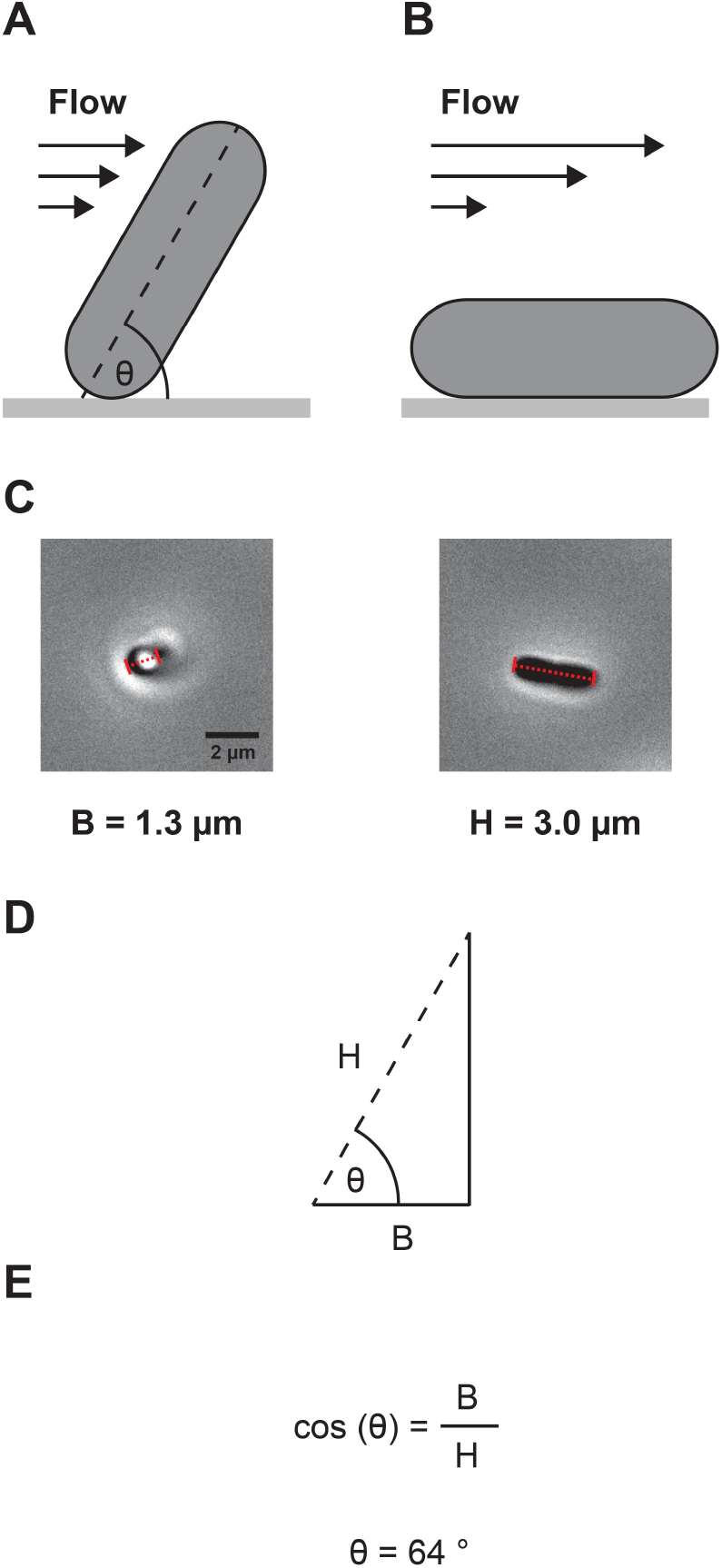
Calculating the cell-surface angle of an example *P. aeruginosa* cell **(A)** A representative *P. aeruginosa* cell in experimental flow condition. The cell has a cell-surface angle θ. **(B)** The same cell exposed to a flow rate of 100 µL/min (shear rate of 8,000 s^-1^). The cell is pushed to the surface and has no cell-surface angle. **(C)** Phase-contrast images of a *P. aeruginosa* cell. The first image is the cell in experimental flow condition. The cell has an observed cell length of 1.3 µm. The second image is of the cell in 100 µL/min (shear rate of 8,000 s^-1^) of flow pushed flat against the surface. The cell has a true cell length of 3.0 µm. **(D)** A right-angle triangle formed by a *P. aeruginosa* cell with a cell-surface angle. The base (B) of the triangle is the observed cell length of the tilted cell in the experimental flow condition under the microscope. The hypothenuse of the triangle (H) is the true cell length in 100 µL/min (shear rate of 8,000 s^-1^) of flow. **(E)** Trigonometry equation used to calculate the cell-surface angle θ. Using the measured observed cell length and true cell length, the cell-surface angle for this example cell was calculated to be ∼64°.

**Figure S7:**
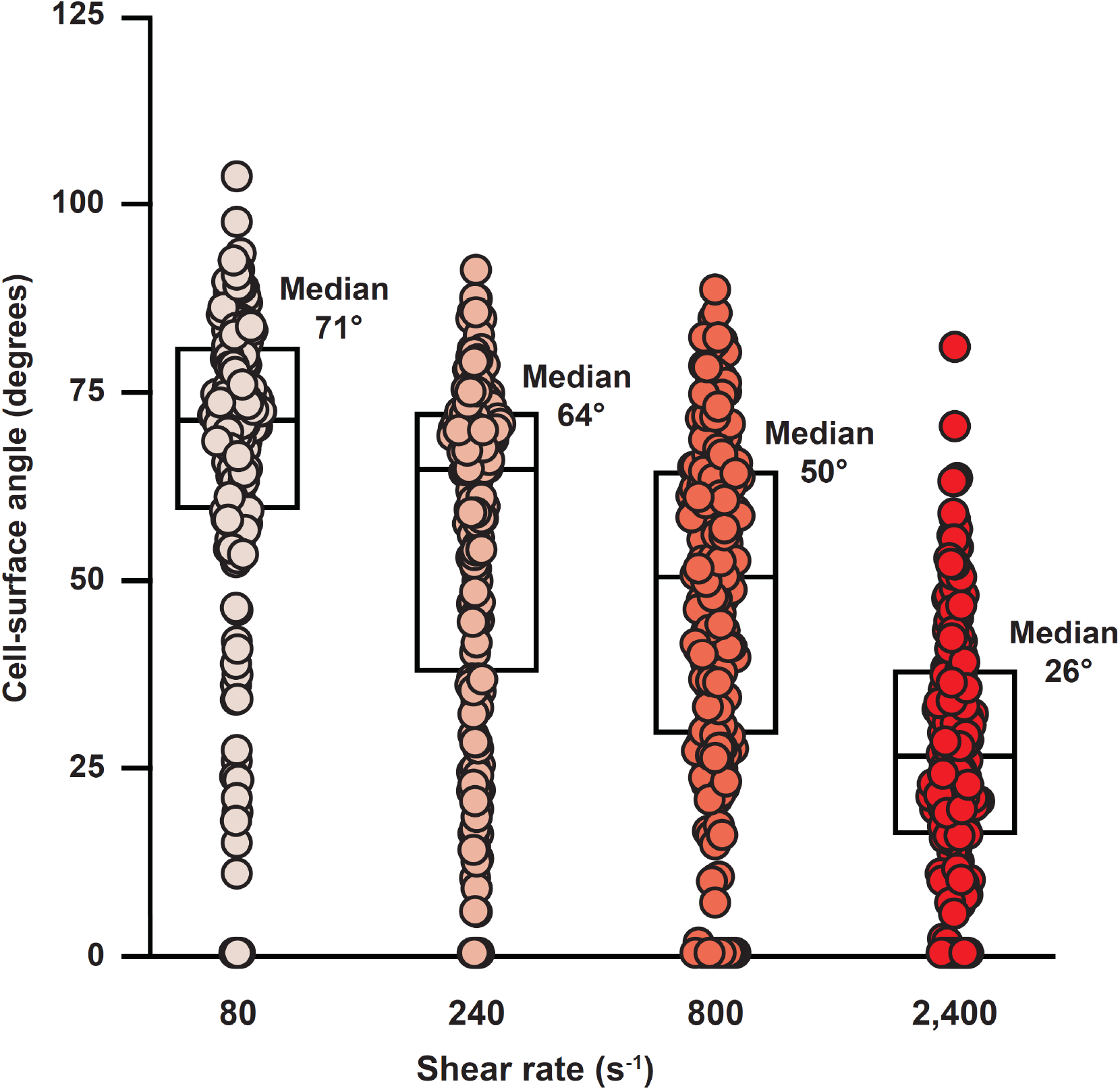
Increasing flow results in a decrease in cell-surface angle of individual cells Quantification of the cell-surface angle of individual WT cells exposed to varied shear rates. N = 180 cells for each shear rate. Statistical significance was determined using Welch’s one-way ANOVA followed by Dunnett’s T3 test. All four experimental conditions were statistically different from one another (*p* < 0.01). Boxplot represents the 25th percentile, median, and 75th percentile for each condition.

